# Perspectives on Quorum sensing in Fungi

**DOI:** 10.1101/019034

**Authors:** Sarangam Majumdar, Subhoshmita Mondal

## Abstract

Keeping a pace with Quorum sensing, analyzing communication shows the close co-evolution of fungi with organisms present in their environment giving insights into multispecies communication. Subsequently, many examples of cell density dependent regulation by extracellular factors have been found in diverse microorganisms. The widespread incidence of diverse quorum-sensing systems strongly suggests that regulation in accordance with cell density is important for the success of microbes in many environments. The paper includes the basic autoregulatory quorum sensing molecules that has been perceived. Although fungal quorum sensing research is still in its infancy, its discovery has changed our views about the fungal kingdom and could eventually lead to the development of new antifungal therapeutics.

## INTRODUCTION

Innumerable variety of fungal organisms represents a major challenge when establishing a homogeneous designation of the signaling processes employed. The population level regulation, also known as “Quorum sensing” has enabled bacterial cells to chemically measure the density of the surrounding population [1]. Subsequently, many examples of cell density dependent regulation by extracellular factors have been found in diverse fungi family. It has become apparent that fungi, like bacteria, also use quorum regulation to affect population-level behaviors such as biofilm formation and pathogenesis [2]. Since the nature and mode of action of communication is as diverse as the response to the information it carries. Inter- and intra species communication has been widely studied analyzing the exchange of information between fungal species [3].The existence of fungal quorum sensing systems reveal that alcohols (farnesol, tyrosol, phenylethanol and tryptophol), peptides (pheromones), lipids (oxylipins), acetaldehydes and few volatile gas and compounds are found to act as quorum sensing molecules regulating morphogenesis, filmentation, pathogenesis as well as various other functions. As described and reviewed, quorum quenching systems of genus such as *Aspergillus, Candida, Saccharomyces* and others has been identified to modulate fungal signaling pathways which are the sensors in cell culture density and biofilm formation. Because of the role of quorum-sensing regulation in biofilm formation, there is particular interest in identifying chemical agents that can control biofilm-associated infections and attenuate chronic infections that involve a biofilm-like state [4]. In the same way that pathogens appear to use quorum-sensing regulation to control the production of virulence determinants, quorum-sensing pathways may have roles within protective microbial communities. However, in most cases the actual molecular mechanism of such communication remains for most parts unknown. The determination of these pathways is of substantial significance as molecular messengers control the expression of fungal virulence determinants including the hyphal switch and biofilm formation or mycotoxin formation. An important issue in the field is the absence of a consensus regarding the definition of fungal QSM that suggested and proposed fungal QSM should:(1) accumulate in the extracellular environment during fungal growth; (2) accumulate in a concentration that is proportional to the population cell density with its effects restricted to a specific stage of growth; (3) induce a coordinated response in the entire population that is not simply an adaptation meant to metabolize or detoxify the molecule itself after a threshold concentration is reached;(4) not be solely a byproduct of fungal catabolism [5]. There are only a few fungal QSM isolated so far where different solutions for intercellular communication during the immense evolutionary gap could have developed, these criteria might need to be refined in the future. Once the importance of quorum sensing is established in pathogenic fungi and the mechanistic details are uncovered, the value of quorum-sensing pathways as potential therapeutic targets can be assessed.

## ELEMENTS OF QUORUM-SENSING REGULATION

In both bacteria and fungi, quorum sensing is mediated by small diffusible signaling molecules that accumulate in the extracellular environment. The signal molecules themselves are generally, though not always, specific to a species or strain, and there is a high degree of structural diversity among the signaling molecules produced by different microbes [6,7,8]. The mechanism by which a signal accumulates in the medium depends on the system; signal export has been shown to involve passive diffusion across the membrane, the action of efflux pumps, and specific transporters [9]. When a signal accumulates to a sufficiently high concentration, the cognate response regulator is activated within the local population of cells, leading to coordinated gene expression. While the details of signal detection in fungi have yet to be elucidated, a number of different strategies for signal detection in bacteria have been described, including cytoplasmic, signal binding transcription factors and cell-surface localized receptors [10]. A number of recent reviews provide an overview of the literature describing fungi have been found to produce extracellular molecules that modulate cellular morphology. In addition to quorum-sensing molecules that affect morphology in fungi, autoregulatory molecules with effects on growth have also been observed.

### Peptides: Pheromones

Since 1959, pheromones have been known to act as an informative molecule and were reported to be involved in the sexual cycle of fungi in 1974 [11]. In the fungal kingdom, they are involved in the reconnaissance of compatible sexual partner to promote plasmogamy and karyogamy between two opposite mating types followed by meiosis. Taking the example of the extensively described sexual cycle of *Saccharomyces cerevisiae*, pheromones are diffusible peptides called a-factor (12 aa). Each mating type responds to the opposite factor, and is able to produce only one of the two peptide pheromones depending on the alleles present at the *MAT* locus [12]. *MFα1* encodes the pheromone precursor, prepro-*α*-factor, which undergoes several proteolytic reactions in the classical secretory pathway before releasing the mature pheromone [13]. Once released, pheromones freely diffuse in the environment and create a concentration gradient. Phenotypically, the morphological response of cells to opposite mating pheromone is the development of a shmoo, that is, directional cell growth in response to the pheromone gradient. As each opposite cell develops a shmoo, the plasmogamy between the two cells occurs when the shmoos establish contact, starting the first step of the sexual cycle. Moreover, *Candida albicans* has not yet been described to undergo meiosis. This yeast displays only an imperfect sexual cycle, where karyogamy results in the formation of tetraploid cells that restore natural diploidy via random loss of chromosomes [14]. This process occurs only after *C. albicans* has undergone a so-called white-to opaque switch where opaque cells are the sole mating competent form of this yeast. The opaque cells morphologically respond to pheromone by producing a shmoo, like in *S. cerevisiae*, but the mating incompetent form, white cells, is also sensitive to pheromone [15, 16]. Indeed, *C. albicans α*-factor but also a-factor promotes the formation of biofilm by white cells via enhancing their adhesiveness. The formation of fungal biomass by white cells facilitates the establishment of a pheromone gradient in a population of individual cells and assists the mating process of opaque cells. This process involves another molecule, farnesol, as the production of this molecule under aerobic conditions induces the death of the mating competent opaque cells. Anaerobic conditions that prevent production of farnesol facilitate mating between *C. albicans* opaque cells. The mechanism of pheromone communication has a broad significance in diverse fungi including ascomycetes like *Histoplasma capsulatum* or *Aspergillus fumigatus*, to basidiomycetes such as *Cryptococcus neoformans* and *Ustilago maydis*, which possess a tetrapolar mating system [17]. Pheromone communication appears to be a critical mechanism for fungi as it supports the exchange of genetic material between cells and by extension the ability of the organism to evolve in response to their environment.

### Lipids: Oxylipins

Oxylipins are oxygenated fatty acids used as cell messengers and have been intensely studied in plants and mammalian cells [18]. They also appear to be widely synthesized and secreted by fungi. *A. nidulans* was reported to produce one of the first oxylipins called psi factor (precocious sexual inducer), which is composed of a series of different oxylipin derivates from oleic acid, linoleic acid, and linolenic acid. The genes involved in the production of psi factor are called *Ppo*s (for psi-producing oxygenases) [19, 20]. Inactivation of these genes results in perturbations not only of psi factor production but also mycotoxins production, as well as in the ratio between the development of sexual and asexual ascospores. Overexpression or addition of psiB*α* or psiC*α* to the culture medium stimulates sexual sporulation and represses asexual spore development while an opposite effect is observed for psiA*α* and psiB*β* [21]. Secondary metabolite mycotoxin sterigmatocystin (ST) and antibiotic penicillin (PN) production are also dependent on oxylipin [22]. Finally, using confocal laser scanning microscopy, oxylipins have been described to accumulate in the capsule of *C. neoformans* before being released into the external medium under the form of hydrophobic droplets that are transported via tubular protuberances [23]. Nigam et al. have described a 3(*R*)-Hydroxytetradecanoic acid, a derivate of linoleic acid as a novel QSM of *C. albicans* [24]. Previously it was known to be produced during the sexual phase of *Dipodascopsis uninucleata*, this oxylipin increases filamentation in *C. albicans* in response to N-acetylglucosamine [25].

### Volatile Compounds and Gas

In addition to releasing mediators into solution or onto solid growth media, organisms also exchange information via the liberation of messenger molecules into air. In the fungal kingdom, as early as in the 1970s, volatile compounds from fungi and others organism have been described to impact on fungal growth [26, 27]. It has been observed that *S. cerevisiae* colonies grown on complex agar form a turbid path in the vicinity of another colony. Subsequently, they discovered that this reaction is induced by the small volatile messenger molecule, later described as ammonia [28], which also required aminoacid uptake for its production. *Trichoderma* species have been described to produce the volatile molecule 6-Pentyl-*α*-pyrone, a secondary metabolite with antifungal activity [29]. However, the induction of conidiation in *Trichoderma* species, which is known to be regulated by a circadian cycle, has also been shown to be controlled via a volatile agent. 8-carbon compounds molecules 1-octen-3-ol, 3-octanol and 3-octanone specifically induce conidiation in colonies placed in the dark. This regulation could involve a calcium-dependant signaling pathway [30]. Fungi are not only responsive to volatile compounds that they produce but also, in at least one example, to a gas liberated during respiration: carbon dioxide (CO_2_). It was demonstrated that the optimum CO_2_ concentration for the germination of *Aspergillus niger* conidiospores is reached not under normal atmospheric concentrations of CO_2_ (0.033%) but at 0.5% [31]. Since then several additional phenotypes in fungi have been attributed to changes in the concentration of environmental CO_2_ including the sporulation of *Alternaria crassa* and *Alternaria cassiae*, conidiation of *Neurospora crassa*, or capsule formation and mating in *C. neoformans* [32]. The significant advances have been made in the understanding of CO_2_ sensing in fungi. It was already known that the yeast to hyphae morphological switch in *C. albicans* is triggered by elevated environmental CO_2_ [33].

### Acetaldehyde

Acetaldehyde, an organic compound involved in several cellular pathways, has been shown to impact on cell-density-dependent glycolytic oscillations of *S. cerevisiae* [34]. In 1964, Chance et al. described that the level of NADH in yeast oscillated when starved cells endure a pulse of glucose after a switch to anaerobic conditions [35]. Since then other metabolites have been described to oscillate in yeast including glucose-6-phosphate, fructose-6-phosphate, fructose-1,6-biphosphate, AMP, ADP, and ATP [36]. Interestingly, at a population level these oscillations are not chaotic but appear to be subject to synchronization. Acetaldehyde was identified as the active molecule in the synchronization of these oscillations, as the use of acetaldehyde traps induced the oscillation to be damped and addition of acetaldehyde to the medium produced a phase shift in the oscillation [34]. Acetaldehyde is a small molecule that can passively diffuse through the cell membrane. No specific target for acetaldehyde is known; however, this compound has an important impact on the NAD+/NADH balance [37]. Acetaldehyde is also a volatile molecule, a property used to study inter and intraspecies communication in a synthetic ecosystem [95]. By engineering sender cells that liberate volatile acetaldehyde and receiver cells that contain a construct under an acetaldehyde-inducible promoter, it was possible to study volatile cell communication in a controlled environment. For example, yeast (*S. cerevisiae*), in combination of sender/receiver for inter and intraspecies resulted in a positive communication between cells [38]. These results show that virtually all cells can communicate with themselves or different species. Evidently such models could bring new insight in the understanding of communication in complex living systems.

### Farnesol

*C. albicans* is a polymorphic fungus normally found in the human microbiota that can cause devastating infections in situations when its relationship with the human host is altered by immune suppression or compromise of epithelial barriers. Polymorphism between yeast, hyphae and pseudohyphae forms is critical for its virulence and corresponds to an adaptive response to environmental changes [39]. Additionally, it has long been observed by experiments that at densities lower than106 cells/ml, *C. albicans* cells develop into filamentous forms, whereas at higher densities the fungus grows as budding yeasts. Hornby and collaborators discovered that this behavior was controlled by a QSM, which they identified as being the isoprenoid farnesol [40]. In the same year, it was described in a different strain of *C. albicans* that farnesoic acid acts like an autoregulatory substance (ARS) also inhibiting filamentation [41]. Farnesoic acid was found only in a strain of *C. albicans* that does not produce farnesol and its effects are less intense. Long before farnesol and farnesoic acid were isolated, a morphogenic autoregulatory substance (MARS) was isolated by Hazen and Cutler that produced similar effects on filamentation of *C. albicans*. However, the chemical features of this molecule differed from farnesol and farnesoic acid and its identity has not been determined yet [42, 43].

### Tyrosol

Tyrosol concentration correlates with the increase of biomass of both planktonic cells and biofilms of *C. albicans* and the addition of tyrosol at early stages of biofilm formation stimulated hyphal growth. It decreases the length of the lag phase of growth, and stimulating filamentation and biofilm formation. These effects are suppressed in the presence of farnesol, suggesting a fine QS-mediated control [44, 45]. Additionally, tyrosol was shown to have an inhibitory activity against neutrophils possibly by interfering with the oxidative burst of these phagocytes [46].

Lastly, the above investigations concern with cell adhesion, pheromone response, calcium/calmodulin, cell integrity, osmotic growth, stress response or cell growth. The interactional context determines the semantic relationship, i.e., the function of the chemical components forms a signal-mediated communication pattern in fungi. Thus, fungal organisms coordinate all their behavioural patterns with a core set of chemical molecules. The interactional context and the different modes of coordinating appropriate response behaviour in development, growth, mating, attack, defense, virulence, etc. determines the combinations of signals that generate the appropriate meaning of function, carrying informational messages.

### Mathematical model

In the other hand, we can also propose stable linear and nonlinear mathematical models of quorum sensing in fungi with the help of partial differential equations and ordinary differential equations. Then stability analysis and the asymptotic analysis can be found out. However, the other factors also seem to come into play and hence statistical methods are needed. Statistical modelling and computational methods can be also taken into account to investigate these density dependent phenomena. Firstly, the statistical variation in morphology throughout a population should be well described. Secondly, a further random phenomenon occurs when the quorum sensing takes place, which is closely linked with the biofilm formation and pathogenesis. Fluid mechanics theory in combination with statistical models will dictate the mathematical equations that need to be solved. A key role in continuum mechanics is played by the concept of representative volume element (RVE). Technically speaking, this requires erotic, stationary random fields of microscale properties and what results is a macroscopic homogeneous continuum. More specifically, we consider finite “windows of observation” which we increase in size.The windows represent statistical volume elements and provide a heterofore missing link to stochastic finite element methods. Further studies may include the use of finite element software in combination with experimental validation studies.

## References

[1] Fuqua, W. C., S. C. Winans, and E. P. Greenberg. 1994. Quorum sensing in bacteria: the LuxR-LuxI family of cell density-responsive transcriptional regulators. J. Bacteriol. 176:269–275.

[2] Reynolds, T. B., and G. R. Fink. 2001. Bakers’ yeast, a model for fungal biofilm formation. Science 291:878–881.

[3] M. T. Tarkka, A. Sarniguet, and P. Frey-Klett, “Inter-kingdom encounters: recent advances in molecular bacterium-fungus interactions,” Current Genetics, vol. 55, no. 3, pp. 233–243, 2009.

[4] Rice, S. A., D. McDougald, N. Kumar, and S. Kjelleberg. 2005. The use of quorum-sensing blockers as therapeutic agents for the control of biofilmassociated infections. Curr.Opin.Investig. Drugs 6:178–184.

[5] Raina S, De Vizio D, Odell M, et al. Microbial quorum sensing: a tool or a target for antimicrobial therapy? Biotech Appl Biochem 2009; 54:65 – 84.

[6] Federle, M. J., and B. L. Bassler. 2003. Interspecies communication in bacteria. J. Clin. Investig. 112:1291–1299.

[7] Henke, J. M., and B. L. Bassler. 2004. Bacterial social engagements. Trends Cell Biol. 14:648–656.

[8] Visick, K. L., and C. Fuqua. 2005. Decoding microbial chatter: cell-cell communication in bacteria. J. Bacteriol. 187:5507–5519.

[9] Pearson, J. P., C. Van Delden, and B. H. Iglewski. 1999. Active efflux and diffusion are involved in transport of Pseudomonas aeruginosa cell-to-cell signals. J. Bacteriol. 181:1203–1210.

[10] Hui, F. M., and D. A. Morrison. 1991. Genetic transformation in *Streptococcus pneumoniae*: nucleotide sequence analysis shows comA, a gene required for competence induction, to be a member of the bacterial ATPdependent transport protein family. J. Bacteriol. 173:372–381.

[11] L. M. Hereford and L. H. Hartwell, “Sequential gene function in the initiation of Saccharomyces cerevisiae DNA synthesis,” Journal of Molecular Biology, vol. 84, no. 3, pp. 445–461, 1974.

[12] J. E. Haber, “Mating-type gene switching in Saccharomyces cerevisiae,” Annual Review of Genetics, vol. 32, pp. 561–599, 1998.

[13] J. P. McGrath and A. Varshavsky, “The yeast STE6 gene encodes a homologue of the mammalian multidrug resistance P-glycoprotein,” Nature, vol. 340, no. 6232, pp. 400–404, 1989.

[14] R. J. Bennett and A. D. Johnson, “Completion of a parasexual cycle in Candida albicans by induced chromosome loss in tetraploid strains,” EMBO Journal, vol. 22, no. 10, pp. 2505– 2515, 2003.

[15] D. R. Soll, “Why does Candida albicans switch?” FEMS Yeast Research, vol. 9, no. 7, pp. 973–989, 2009.

[16] K. J. Daniels, T. Srikantha, S. R. Lockhart, C. Pujol, and D. R. Soll, “Opaque cells signal white cells to form biofilms in Candida albicans,” EMBO Journal, vol. 25, no. 10, pp. 2240–2252, 2006.

[17] G. Bakkeren, J. Kämper, and J. Schirawski, “Sex in smut fungi: structure, function and evolution of mating-type complexes,” Fungal Genetics and Biology, vol. 45, supplement 1, pp. S15–S21, 2008.

[18] A. Mosblech, I. Feussner, and I. Heilmann, “Oxylipins: structurally diverse metabolites from fatty acid oxidation,” Plant Physiology and Biochemistry, vol. 47, no. 6, pp. 511–517, 2009.

[19] A. M. Calvo, L. L. Hinze, H. W. Gardner, and N. P. Keller, “Sporogenic effect of polyunsaturated fatty acids on development of Aspergillus spp,” Applied and Environmental Microbiology, vol. 65, no. 8, pp. 3668–3673, 1999.

[20] D. I. Tsitsigiannis, T. M. Kowieski, R. Zarnowski, and N. P. Keller, “Three putative oxylipin biosynthetic genes integrate sexual and asexual development in Aspergillus nidulans,” Microbiology, vol. 151, no. 6, pp. 1809–1821, 2005.

[21] D. I. Tsitsigiannis, R. Zarnowski, and N. P. Keller, “The lipid body protein, PpoA, coordinates sexual and asexual sporulation in Aspergillus nidulans,” Journal of Biological Chemistry, vol. 279, no. 12, pp. 11344–11353, 2004.

[22] D. I. Tsitsigiannis and N. P. Keller, “Oxylipins act as determinants of natural product biosynthesis and seed colonization in Aspergillus nidulans,” Molecular Microbiology, vol. 59, no. 3, pp. 882–892, 2006.

[23] O. M. Sebolai, C. H. Pohl, P. J. Botes et al., “3-hydroxy fatty acids found in capsules of *Cryptococcus neoformans*,” Canadian Journal of Microbiology, vol. 53, no. 6, pp. 809–812, 2007.

[24] S. Nigam, R. Ciccoli, I. Ivanov, M. Sczepanski, and R. Deva, “On mechanism of quorum sensing in Candida albicans by 3(R)-hydroxy-tetradecaenoic acid,” Current Microbiology, vol. 62, no. 1, pp. 55–63, 2011.

[25] P. Venter, J. L. F. Kock, G. Sravan Kumar et al., “Production of 3R-hydroxy-polyenoic fatty acids by the yeast *Dipodascopsis uninucleata*,” Lipids, vol. 32, no. 12, pp. 1277–1283, 1997.

[26] N. Fries, “Effects of volatile organic compounds on the growth and development of fungi,” Transactions of the British Mycological Society, vol. 60, pp. 1–21, 1973.

[27] S. A. Hutchinson, “Biological activities of volatile fungal metabolites,” Annual Reviews of Phytopathology, vol. 11, pp. 223–246, 1973.

[28] Z. Palkova, B. Janderova, J. Gabriel, B. Zikanova, M. Pospisek, and J. Forstova, “Ammonia mediates communication between yeast colonies,” Nature, vol. 390, no. 6659, pp. 532–536, 1997.

[29] J. M. Cooney and D. R. Lauren, “Trichoderma/pathogen interactions: measurement of antagonistic chemicals produced at the antagonist/pathogen interface using a tubular bioassay,” Letters in Applied Microbiology, vol. 27, no. 5, pp. 283–286, 1998.

[30] G. Muthukumar, E. C. Jensen, A. W. Nickerson, M. K. Eckles, and K. W. Nickerson, “Photomorphogenesis in Penicillium isariaeforme: exogenous calcium substitytes for light,” Photochemistry and Photobiology, vol. 53, no. 2, pp. 297–291, 1991.

[31] J. R. Vakil, M. R. Raghavendra Rao, and P. K. Bhattacharyya, “Effect of CO_2_ on the germination of conidiospores of *Aspergillus niger*,” Archiv für Mikrobiologie, vol. 39, no. 1, pp. 53–57, 1961.

[32] Y. S. Bahn, G. M. Cox, J. R. Perfect, and J. Heitman, “Carbonic anhydrase and CO_2_ sensing during ***Cryptococcus neoformans* growth, differentiation, and virulence,”** Current Biology, vol. 15, no. 22, pp. 2013–2020, 2005.

[33] W. Sims, “Effect of carbon dioxide on the growth and form of *Candida albicans*,” Journal of MedicalMicrobiology, vol. 22, no. 3, pp. 203–208, 1986.

[34] P. Richard, B. M. Bakker, B. Teusink, K. Van Dam, and H. V. Westerhoff, “Acetaldehyde mediates the synchronization of sustained glycolytic oscillations in populations of yeast cells,” European Journal of Biochemistry, vol. 235, no. 1–2, pp. 238– 241, 1996.

[35] B. Chance, R. W. Estabrook, and A. Ghosh, “Damped sinusoidal oscillations of cytoplasmic reduced pyridine nucleotide in yeast cells,” Proceedings of the National Academy of Sciences of the United States of America, vol. 51, pp. 1244–1251, 1964.

[36] P. Richard, “The rhythm of yeast,” FEMSMicrobiology Reviews, vol. 27, no. 4, pp. 547–557, 2003.

[37] A. Betz and J. U. Becker, “Phase dependent phase shifts induced by pyruvate and acetaldehyde in oscillating NADH of yeast cells,” Journal of Interdisciplinary Cycle Research, vol. 6, no. 2, pp. 167–173, 1975.

[38] W. Weber, M. Daoud-El Baba, and M. Fussenegger, “Synthetic ecosystems based on airborne inter-and intrakingdom communication,” Proceedings of the National Academy of Sciences of the United States of America, vol. 104, no. 25, pp. 10435–10440, 2007.

[39] Gow NA, Brown AJ, Odds FC. Fungal morphogenesis and host invasion. Curr Opin Microbiol 2002; 5: 366 – 371.

[40] Hornby JM, Jensen EC, Lisec AD, et al. Quorum sensing in the dimorphic fungus *Candida albicans* is mediated by farnesol. Appl Environ Microbiol 2001; 67 : 2982 – 2992.

[41] Oh KB, Miyazawa H, Naito T, Matsuoka H. Purification and characterization of an autoregulatory substance capable of regulating the morphological transition in ***Candida albicans***. Proc Natl Acad Sci USA 2001; 98 : 4664 – 4668.

[42] Nickerson KW, Atkin AL, Hornby JM. Quorum sensing in dimorphic fungi: farnesol and beyond. Appl Environ Microbiol 2006; 72 : 3805 – 3813.

[43] Hazen KC, Cutler JE. Isolation and purification of morphogenic autoregulatory substance produced by ***Candida albicans***. J Biochem 1983; 94 : 777 – 783.

[44] Chen H, Fujita M, Feng Q, Clardy J, Fink GR. Tyrosol is a quorumsensing molecule in *Candida albicans*. Proc Natl Acad Sci USA 2004; 101: 5048 – 5052.

[45] Alem MA, Oteef MD, Flowers TH, Douglas LJ. Production of tyrosol by *Candida albicans* biofilms and its role in quorum sensing and biofilm development. Eukaryot Cell 2006; 5 : 1770 – 1779.

[46] Cremer J, Vatou V, Braveny I. 2,4-[hydroxyphenyl]-ethanol, an antioxidative agent produced by *Candida* spp., impairs neutrophilic yeast killing in vitro. FEMS Microbiol Lett 1999; 170 : 319 –325.

